# Nicotine in E-Cigarette Aerosol Reduces GABA Neuron Migration via the α7 Nicotinic Acetylcholine Receptor

**DOI:** 10.1101/2025.09.11.675617

**Authors:** Amber A. Parnell, Deirdre M. McCarthy, Mia X. Trupiano, Christopher Schatschneider, Pradeep G. Bhide

**Affiliations:** Department of Biomedical Sciences, Florida State University College of Medicine, Tallahassee, FL; Program in Neuroscience, Florida State University, Tallahassee, FL; Institute for Pediatric Rare Diseases, Florida State University College of Medicine, Tallahassee, FL; Department of Psychology, College of Arts and Sciences, Florida State University, Tallahassee, FL, United States

**Author notes:** **Corresponding Author:** Pradeep G. Bhide, Ph.D., Department of Biomedical Sciences, FSU College of Medicine, 1115, West Call, Street, Tallahassee, FL 32306.

**Keywords:** ganglionic eminence, GAD67-GFP, e-liquid, interneuron, brain development, prenatal nicotine

## Abstract

Prenatal nicotine exposure is linked to adverse neurodevelopmental outcomes, yet e-cigarette use during pregnancy continues to rise due to aggressive marketing efforts and misconceptions of safety. We investigated the effect of prenatal e-cigarette aerosol exposure on the migration of GABA neurons, a developmental process critical for the establishment of cerebral cortical circuitry. Pregnant mice were exposed to nicotine-containing aerosol (e-cigarette), nicotine-free aerosol (e-liquid) or room air (control) daily beginning 2 weeks before conception and continuing until gestational day 14. E-cigarette, but not e-liquid, aerosol significantly reduced GABA neuron density in the dorsal cerebral wall at rostral forebrain level and within the marginal zone, reflecting region-specific vulnerabilities. *In vitro* explant cultures revealed that nicotine dose-dependently reduced neuronal migration, and this effect was mimicked by a selective α7 nicotinic acetylcholine receptor (nAChR) agonist. Blocking the α7 nAChR using a selective antagonist attenuated the effects of nicotine on neuronal migration. These findings reveal a previously unrecognized vulnerability of GABA neuron migration to e-cigarette aerosol and identify α7 nAChR activation as a mechanism for nicotine-induced impairment of GABA neuron migration. Moreover, the findings highlight the need for translational efforts to update clinical guidance and public policy regarding e-cigarette use during pregnancy.

## Introduction

Electronic cigarettes (e-cigarettes) are battery-operated devices that deliver aerosolized nicotine, often mixed with flavoring agents and other additives. Despite the growing evidence of harm, e-cigarettes are widely perceived by consumers and some healthcare providers as safer alternatives to combustible cigarettes, and as a tool for smoking cessation ^1–3^. These perceptions alongside aggressive marketing strategies, have contributed to a substantial rise in e-cigarette use globally. In the United States, adult e-cigarette use increased from 4.5% in 2019 to 6.5% in 2023 ^4^. Usage among pregnant women is also rising, with an estimated 7–10% reporting use during or around the time of pregnancy ^5–9^.

E-cigarette aerosols expose individuals to not only nicotine but also to a variety of potentially toxic substances, including heavy metals, carbon monoxide, formaldehyde, acetaldehyde, and acrolein— byproducts generated from the aerosolization of e-liquids that are typically composed of propylene glycol and vegetable glycerin ^10–13^.

Extensive clinical evidence indicates that prenatal exposure to nicotine, particularly through combustible cigarette smoking, is associated with adverse pregnancy outcomes such as miscarriage, preterm birth, low birth weight, developmental delays, and sudden infant death syndrome ^14–17^. Moreover, nicotine exposure during pregnancy is implicated in long-term neurodevelopmental consequences, including increased risk of attention deficit hyperactivity disorder, conduct disorders, and cognitive impairments in children ^18–20^.

While the effects of combustible cigarette use during pregnancy as well as direct exposure to nicotine (oral, intravenous or subcutaneous) on behavioral, neurochemical and neuroanatomical parameters are well documented in model organisms ^21–25^, the neurodevelopmental impact of prenatal e-cigarette exposure remains poorly understood. Limited clinical data have focused on birth outcomes ^26–29^, while rodent models have begun to reveal behavioral abnormalities such as hyperactivity, memory deficits, and elevated risk-taking behaviors following prenatal e-cigarette or e-liquid exposure ^30–39^. These behavioral phenotypes are often accompanied by molecular alterations, including changes in gene expression, epigenetic modifications, and neuroinflammatory markers within the brain ^30,31,35,36^.

A converging body of literature suggests that disrupted gamma-aminobutyric acid (GABA) signaling is a shared neurobiological factor across many neurodevelopmental disorders some of which are linked to early-life nicotine exposure ^40–42 43,44^. Yet, a direct investigation of the effects of prenatal e-cigarette aerosol exposure or nicotine exposure in any other form on GABA neuron development in the embryonic brain has not been reported to our knowledge.

Therefore, in this study, we sought to examine the impact of prenatal exposure to e-cigarette and e-liquid aerosols in a mouse model on the migration of GABA neurons from the ganglionic eminence (GE) to the dorsal forebrain, the future cerebral cortex. Migration of GABA neurons is a crucial and early developmental process critical for the proper temporal and spatial positioning and integration of GABA neurons within cortical circuits ^45–47^. By comparing the effects of e-cigarette (with nicotine) and e-liquid (nicotine-free) aerosols, we sought to identify the specific contributions of nicotine and non-nicotine constituents of the aerosols to potential changes in GABA neuron migration. Furthermore, we employed an *in vitro* explant culture system to assess whether nicotine’s effects are mediated via the α7 nicotinic acetylcholine receptor (nAChR), a receptor subtype highly expressed in the embryonic forebrain of humans and rodents ^48–51^. Our findings provide novel insights into the receptor-mediated effects of nicotine on GABA neuron migration, and potential neurodevelopmental risk of prenatal nicotine exposure.

## Results

### E-cigarette and E-liquid Aerosol Exposure

Female Swiss Webster mice (6–8 weeks old) were exposed to e-cigarette or e-liquid aerosol daily in a whole-body aerosol exposure system. E-cigarette aerosol was generated from a solution containing 24 mg/mL of nicotine dissolved in a 50:50 propylene glycol: vegetable glycerin mixture. The non-nicotine containing e-liquid aerosol used the same formulation but without nicotine. A separate cohort of age-matched control female mice was exposed to room air. Following approximately 5-days of habituation to the equipment and an additional 2-weeks of acclimation to the exposure conditions, the mice were bred with exposure-naïve GAD67-GFP^+/−^ males. This breeding scheme produced litters containing GAD67-GFP^+/−^ embryos, in which GABA neurons express GFP facilitating their identification in histological sections of the brain by fluorescence microscopy^52–54^. Daily exposures of the pregnant mice continued until the 14^th^ gestational day (GD; Fig. 1). To examine dose-dependency, two parallel sets of exposures were performed, each lasting either 48 or 96 minutes per day. Maternal serum cotinine levels on GD14 were 35.12 ± 5.91 ng/mL and 75.22 ± 13.11 ng/mL, in the 48 min and 96 min cohorts, respectively. The 96 min exposure paradigm produced maternal cotinine levels comparable to those in our previous study ^55^, using an oral nicotine exposure paradigm (via drinking water), which had demonstrated significant deficits in frontal cortical GABA-to-non-GABA neuron ratios in the adult mouse brain. Therefore, the 96 min exposure paradigm was used in the present study. The cotinine levels in the embryonic brain on embryonic day 14 (E14) from the e-cigarette aerosol exposure group were 41.11 ± 10.98 ng/g and 94.38 ± 24.40 ng/g following 48 min and 96 min of exposure, respectively, indicating that cotinine accumulation in the embryonic brain was nicotine dose dependent.

**Figure 1.**
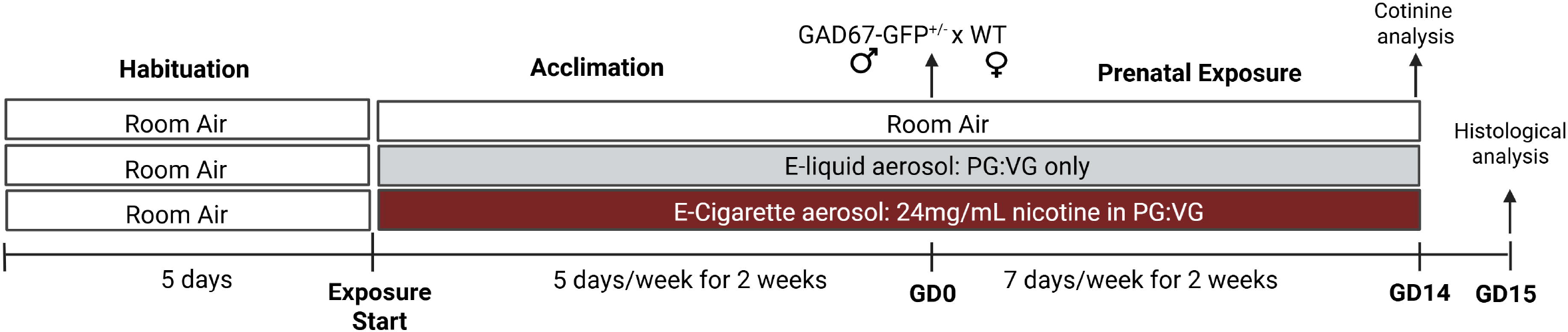
Experimental Design. Female Swiss Webster mice were habituated to the equipment by exposure to room air for 48 or 96 min/day for 5 days. Next, they were acclimated to their exposure conditions (room air, e-liquid or e-cigarette; n=8 per group) daily for 2 weeks. Then, the female mice were bred with unexposed GAD67-GFP^+/−^ heterozygous male mice. Daily exposure of the mice continued through pregnancy until gestational day 14 (GD14; day of conception was designated as GD0). Trunk blood from e-cigarette exposed dams and brains from their embryos were collected on GD14 for cotinine analysis. Brains from GAD67-GFP^+/−^ embryos from each of the three exposure groups were collected on GD15 (24hr. after the final room air or aerosol exposure), for analysis of GABA neuron migration in histological sections of the forebrain.

### Pregnancy and Embryo Metrics

There was no significant difference in the body weights of female mice assigned to the three exposure groups at the time of the assignment or approximately 2 weeks later at the time of conception (Table 1). Since using e-cigarette and other nicotine containing products during pregnancy is associated with adverse pregnancy outcomes and compromised fetal growth ^26–29,55^ we analyzed changes in maternal body weight on the 10^th^, 13^th^, and 15^th^ days of pregnancy as a percentage of body weight on GD0. Although pregnant mice in all groups gained body weight over this period, there was no significant effect of aerosol exposure or a significant interaction between aerosol exposure and gestational day on percent change in body weight, the number of embryos per dam or the crown-rump-length of the embryos at E15 (Table 1). Altogether, aerosol exposures did not produce significant effects on pregnancy or litter metrics.

**Table 1.**
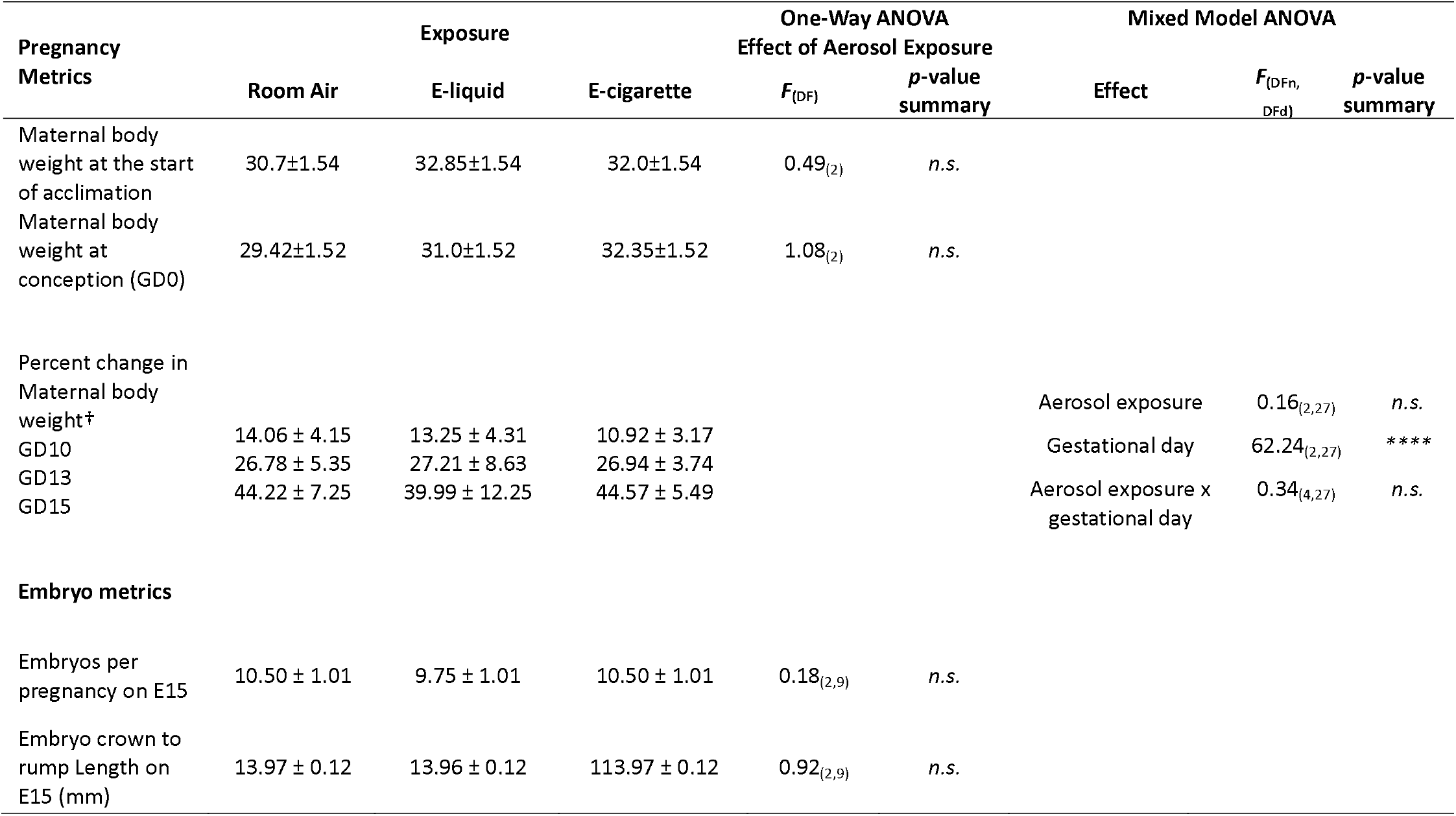
Mean ± SEM values for pregnancy and embryo metrics. † = percentage of baseline bodyweight on gestational day 0 (GD0). E=embryonic day; One-way and Mixed Model ANVOAs. **** *p* < 0.0001; n.s. = not significant.

### GABA Neuron Density in the Dorsal Cerebral Wall

GABA neurons of the forebrain are generated in the ganglionic eminence (GE) of the basal forebrain and migrate to distant forebrain targets, including the cerebral cortex, during the prenatal period ^56,57^. In the embryonic mouse brain, GABA neuron migration from the GE to the dorsal cerebral wall, the future cerebral cortex, begins around embryonic day 11 (E11) and is robust by E15 ^58,59^. To determine if e-cigarette or e-liquid aerosol exposure affected GABA neuron migration, we analyzed the density of GABA neurons in the dorsal cerebral wall by counting the GFP+ cells by fluorescence microscopy in coronal sections of the embryonic brain. In the description below, GFP+ cells are referred to as GABA neurons consistent with previous reports ^52–54^.

The GE, consisting of lateral, medial and caudal subdivisions (LGE, MGE and CGE, respectively), is a rostro-caudally elongated structure characterized by substantial regional heterogeneity in the expression of cell-intrinsic and environmental factors, including transcription factors and extracellular signaling molecules ^60,61^. Moreover, the dorsal cerebral wall exhibits a lateral-to-medial maturational gradient of GABA neurons, such that at any given embryonic day, more mature (“older”) GABA neurons are located laterally, while less mature (“younger”) neurons reside more medially ^58,61,62^. Thus, the basal and the dorsal forebrain exhibit pronounced spatial and temporal heterogeneity, which may influence or modify the effects of aerosol exposures observed in this study. To address this issue, we collected and analyzed GABA neuron density separately at rostral and caudal levels of the forebrain, and at each level, collected data separately at lateral, middle and medial anatomical locations of the dorsal forebrain (Fig. 2A, E). If the aerosol exposures produced significant changes in the density of GABA neurons in the dorsal cerebral wall it would be an indirect indication of changes in the migration of the GABA neurons from the basal to the dorsal forebrain.

**Figure 2.**
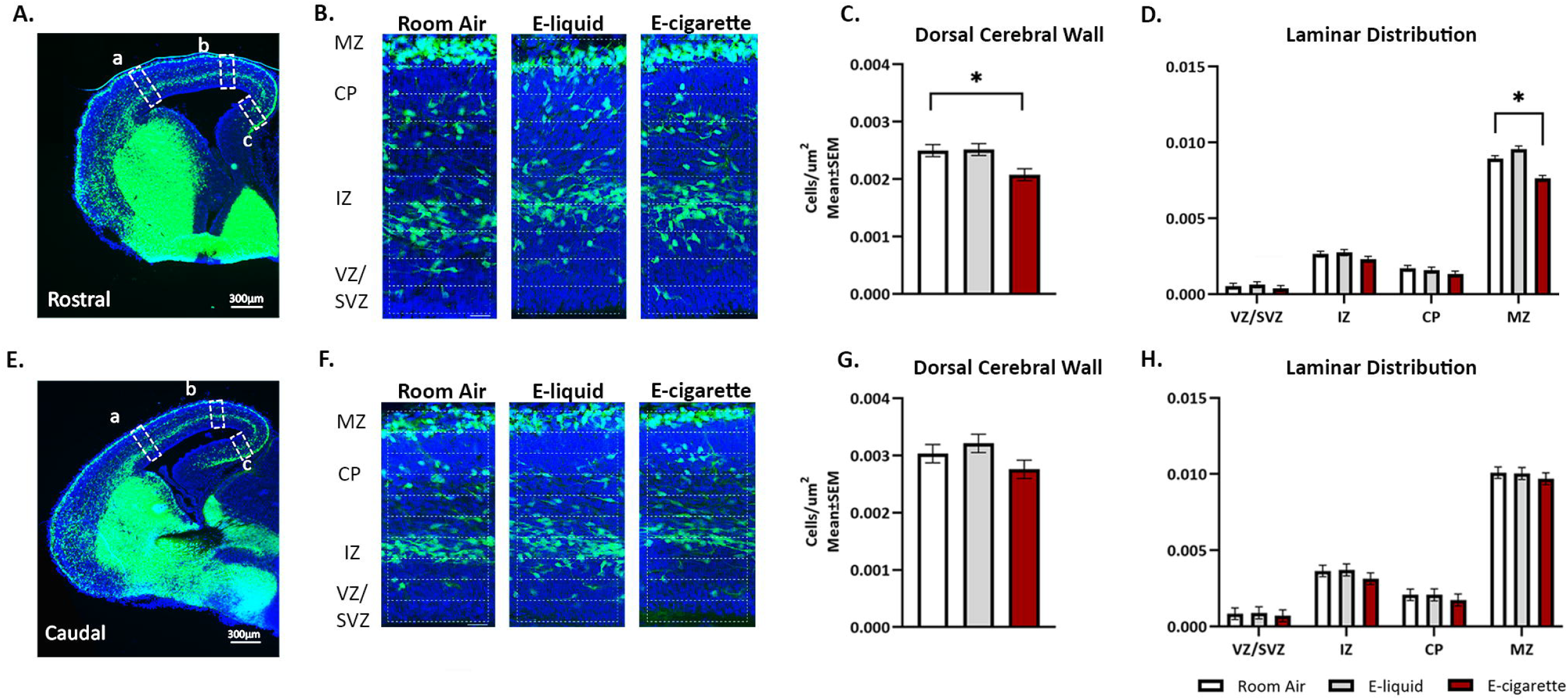
GABA Cell Density in the E15 Dorsal Forebrain. Representative coronal sections at the rostral (**A**) and caudal (**E**) levels of the forebrain from a GAD67-GFP^+/−^ embryo showing GFP+ GABA neurons (green). DAPI (blue) was used to label cell nuclei. GABA neurons were counted within a region of interest (white dashed rectangles) at each of three anatomical locations along the lateral to medial extent of the dorsal cerebral wall: lateral (a), middle (b), and medial (c). A higher magnification view of the “middle” anatomical location (b) of the dorsal cerebral wall from the rostral (B) and caudal (F) levels showing subdivision into 10 equal-sized grids for the purpose of counting GFP+ GABA neurons, and stratification into marginal zone (MZ), cortical plate (CP), intermediate zone (IZ), and subventricular/ventricular zones (SVZ/VZ) (**B, F**). GABA neuron density is reduced in the e-cigarette group compared to the room air and e-liquid groups at the rostral level (**C**) and there is no significant difference among the 3 groups at the caudal level (**G**). GABA neuron density was reduced in the e-cigarette group compared to the room air group in the MZ at the rostral level (**D**) while there were no significant differences among the groups in any lamina at the caudal level (**H**). Scale bars represent 300µm (**A, E**) and 20µm (**B, F).** * = *p* < 0.05 [Pair-wise comparisons with Dunnett’s (**C**) or Benjamini-Hochberg linear step up (**D**) corrections]. Data represent mean±SEM.

### Rostral Forebrain Level

At the rostral level, there was a significant effect of aerosol exposure on GABA neuron density (*F*_(2,54)_ = 5.63, *p* < 0.01). Pair-wise comparisons demonstrated that prenatal e-cigarette aerosol significantly reduced the GABA neuron density compared to room air (Dunnett’s test: *t*_(54)_ = −2.84, *p* < 0.05; Fig. 2C). E-liquid aerosol and room air controls were not significantly different (Dunnett’s test: *t*_(54)_ = 0.89, *p* > 0.05). GABA neuron density significantly differed among anatomical locations (lateral, middle, medial; *F*_(2,54)_ = 7.83, *p* < 0.05). Bonferroni pair-wise comparisons test indicated that the density at the middle location was significantly greater than that at the lateral (*t*_(54)_ = −3.16, *p* < 0.05) and medial (*t*_(54)_ = −3.16, p < 0.05) locations. GABA neuron density was not significantly different between lateral and medial locations (*t*_(54)_ = −0.48, *p* > 0.05). Finally, the interaction between aerosol exposure and anatomical location was not significant (*F*_(4,54)_ = 1.23, *p* > 0.05), suggesting that the reduction in GABA neuron density due to e-cigarette aerosol exposure was not location-specific and occurred throughout the lateral to medial extent of the dorsal cerebral wall.

To determine whether the observed differences in GABA neuron density across the three anatomical locations were attributable to variations in the radial thickness of the dorsal cerebral wall, we analyzed thickness as an independent measure. Anatomical location had a significant effect on radial thickness (*F*_(2,54)_ = 145.85, *p* < 0.0001). Bonferroni multiple comparison tests showed that thickness was significantly greater at the lateral location compared to the middle (*t*_(54)_ = −6.99, *p* < 0.0001), greater at the middle compared to the medial (*t*_(54)_ = 10.00, *p* < 0.0001) and greater at the lateral compared to the medial location (*t*_(54)_ = 16.99, *p* < 0.0001).

We did not detect a significant effect of aerosol exposure on radial thickness at any of the three anatomical locations (*F*_(2,54)_ = 0.57, *p* > 0.05). In addition, there was no significant interaction between aerosol exposure and anatomical location (*F*_(4,54)_ = 0.38, *p* > 0.05) indicating that aerosol exposure did not alter the radial thickness of the dorsal cerebral wall along the lateral-to-medial axis. Therefore, although radial thickness varied by anatomical location, the changes in GABA neuron density observed following e-cigarette aerosol exposure occurred independently of these differences in radial thickness.

GABA neurons migrate through the dorsal cerebral wall in two anatomically distinct zones: the marginal zone (MZ), beneath the pial surface, and the intermediate zone (IZ) located above the subventricular zone (SVZ; Fig 2B, F). The extracellular matrix within each zone presents unique molecular and cellular cues that influence neuronal migration ^61,63^. To assess whether aerosol exposure alters GABA neuron migration in a zone-specific manner, we analyzed GABA neuron density separately in the MZ, IZ, as well as in the SVZ and the intervening cortical plate (CP)—the region where GABA neurons integrate with glutamatergic neurons and ultimately settle into their final laminar positions later in development. The term CP applies only to the lateral and middle locations, which are the primordia of neocortical areas whereas hippocampal plate is the term that applies to the medial location, which is the primordium of the archicortex. The effect of exposure was analyzed in each zone separately within each of the three anatomical locations.

There was a significant effect of aerosol exposure (*F*_(2,242)_ = 8.43, *p* < 0.001), anatomical location (*F*_(2,242)_ = 26.47, *p* < 0.0001), and migratory zone (*F*_(3,242)_ = 1382.62, *p* < 0.0001) on GABA neuron density. There was a significant interaction between exposure and migratory zone (*F*_(6,242)_ = 5.66, *p* < 0.0001; Fig. 2D). Pair-wise comparisons showed that there was a significant reduction in GABA neuron density within the MZ following prenatal e-cigarette aerosol exposure compared to room air (*t*_(242)_ = −4.71, *p* < 0.05).

The interaction between exposure and anatomical location (*F*_(4,242)_ = 1.20, *p* > 0.05) and the three-way interaction between exposure, location, and migratory zone (*F*_(12,242)_ = 0.81, *p* > 0.05) were not significant. Thus, e-cigarette aerosol exposure reduced GABA neuron density selectively in the MZ.

### Caudal Forebrain Level

Our analyses did not show a significant effect of aerosol exposure (*F*_(2,54)_ = 2.02, *p* > 0.05; Fig. 2G) or interaction between aerosol exposure and location (*F*_(4,54)_ = 0.78, *p* > 0.05) on GABA neuron density. There was a significant difference in neuron density among the three anatomical locations (*F*_(2,54)_ = 16.48, *p* < 0.0001). Bonferroni multiple comparison tests indicated that GABA neuron density was greater at the medial location compared to the middle (*t*_(54)_ = −5.28, *p* < 0.0001) and lateral locations (*t*_(242)_ = −4.59, *p* < 0.0001), but it did not significantly differ between lateral and middle locations (*t*_(54)_ = −0.69, *p* > 0.05). There was a significant effect of location on the radial thickness of the caudal cerebral wall (*F*_(2,54)_ = 355.99, *p* < 0.0001). Bonferroni multiple comparison test revealed that the radial thickness of the lateral region was greater than the middle (*t*_(54)_ = −14.22, *p* < 0.0001), middle was greater than the medial (*t*_(54)_ = 12.44, p < 0.0001) and lateral was greater than the medial (*t*_(54)_ = 26.66, *p* < 0.0001). However, the radial thickness was not significantly influenced by the aerosol exposure (*F*_(2,54)_ = 0.69, *p* > 0.05) or the interaction between aerosol exposure and anatomical location (*F*_(4,54)_ = 0.14, *p* > 0.05).

Additionally, the effect of aerosol exposure on GABA neuron density in the different migratory zones was not significant (*F*_(2,242)_ = 1.00, *p* > 0.05). The interactions between exposure and location (*F*_(4,242)_ = 0.84, *p* > 0.05), exposure and migratory zone (*F*_(6,242)_ = 0.25, *p* > 0.05; Fig. 2H), and the three-way interaction between exposure, location, and migratory zone were not significant (*F*_(12,242)_ = 0.56, *p* > 0.05).

Thus, e-cigarette aerosol but not e-liquid aerosol exposure produced significant decreases in GABA neuron density at the rostral level of the forebrain but not at the caudal level. The effects were significant in the MZ and were observed at each of the three anatomical locations across the lateral to medial extent.

### Nicotine reduces GABA Neuron Migration In Vitro

The preceding data demonstrated that prenatal exposure to e-cigarette aerosol, but not e-liquid aerosol, resulted in a reduction in GABA neuron density. As nicotine was the only component distinguishing the two aerosol types, we attributed this effect to nicotine alone. To directly test whether nicotine impairs GABA neuron migration from the basal forebrain, we employed an *in vitro* explant culture approach^52,53^. We sectioned brains from E15 exposure-naïve embryos on a Vibratome and collected rostral forebrain sections corresponding to those chosen to represent the rostral forebrain level in the *in vivo* studies based on anatomical landmarks. Since the *in vivo* results indicated significant effects of e-cigarette aerosol only at the rostral level, we used basal forebrain explants exclusively from sections representing the rostral forebrain level (Fig. 3A). The use of Vibratome sections ensured consistency in explant thickness across brains and allowed us to match the explant location precisely to the *in vivo* sampling site, enabling direct comparison between the *in vivo* and *in vitro* findings. However, neither the *in vivo* nor *in vitro* approaches isolated cells exclusively derived from the medial or lateral ganglionic eminence (MGE or LGE). Therefore, both datasets likely included a mixture of MGE- and LGE-derived cells.

**Figure 3.**
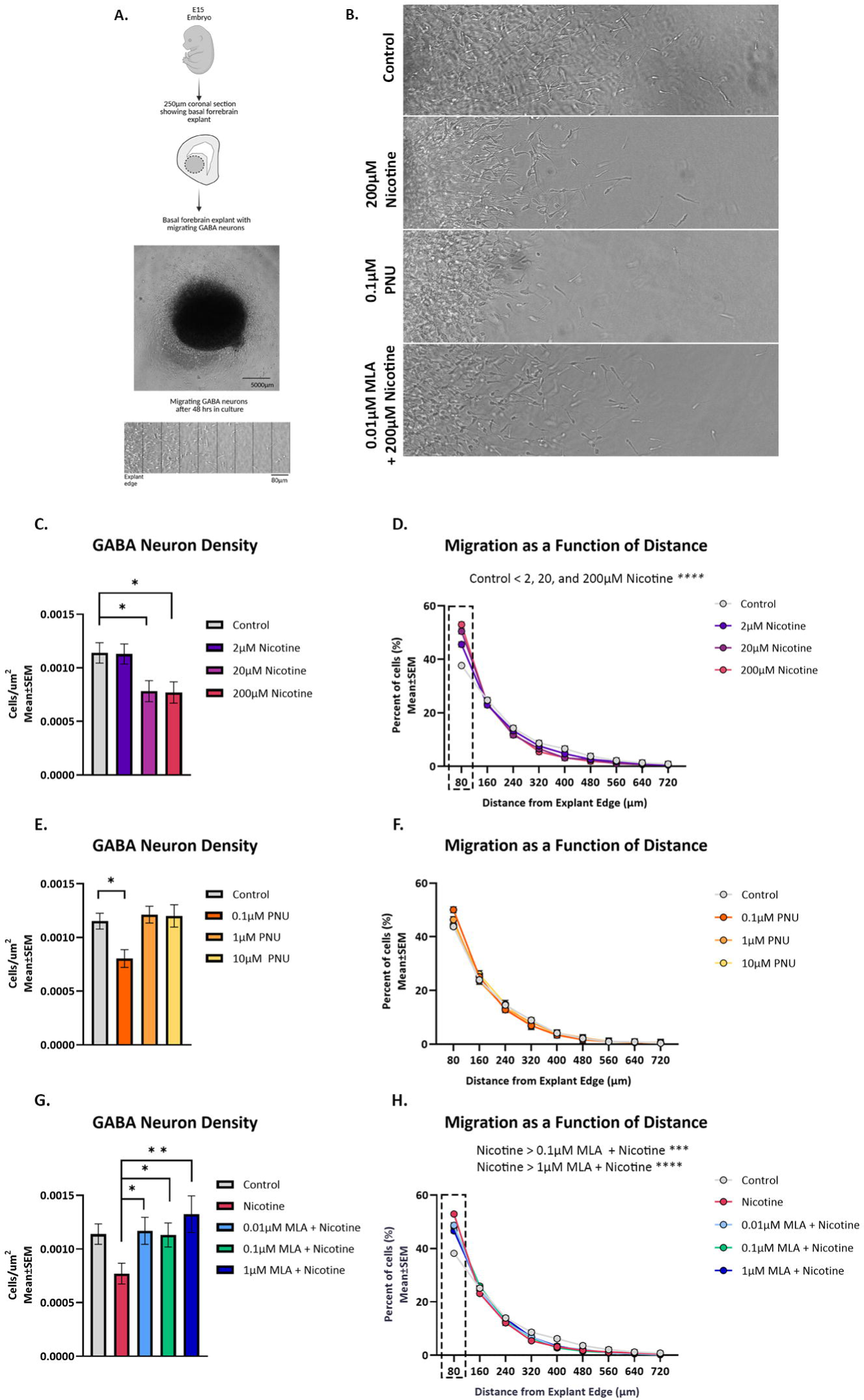
GABA neuron migration in explant cultures. An overview of the in vitro experimental design and outcomes (**A**) showing the principal steps of collection of basal forebrain eminence explants from coronal sections of E15 rostral forebrains, culturing the explants in Matrigel, and migration of GABA neurons from the explants following 48 hr. in culture shown as a cartoon. In addition, a photomicrograph showing neurons migrating away from the explant edge, and the subdivision of the migration field into 80 µm-wide bins for quantitative analysis is shown. Representative micrographs showing neuronal migration under the different drug exposure conditions (B). Quantitative analyses showed that the density of GABA neurons migrating away from GE explants was reduced in the presence of 20µM and 200µM nicotine compared to control, whereas 2µM nicotine did not produce significant changes (**C**). GABA neuron density as a percentage of the total number of migrating GABA neurons increased within the initial 80µm of the explant edge (dashed box) following treatment with nicotine (2, 20 and 200µM; **D**). The density of GABA neurons migrating from the explant was significantly reduced following treatment with the selective α7 nAChR agonist PNU282987 at 0.1µM concentration compared to control (0.01% DMSO), whereas 1µM or 10µM concentrations of PNU282987 did not produce significant differences (**E**). None of the 3 concentrations of PNU282987 significantly altered the percentage of migrating GABA neurons located within the initial 80µm of the explant edge compared to control (**F**). The density of migrating GABA neurons was significantly higher in the explants exposed to the α7 nAChR antagonist methyllycaconitine (MLA; 0.01µM, 0.1µM, 1µM) 2 hr. prior to nicotine exposure compared to the density in the explants exposed to nicotine (200 µM) alone, and not significantly different from the density in the control explants (**G**). For comparative analyses, data from explants treated with 200µM nicotine and control explants (plain media), shown in panel C was re-incorporated. The percentage of cells within the initial 80μm interval (dashed box) in explants treated with MLA 2 hr. prior to nicotine exposure was significantly higher than the percentage in the control explants (0.01, 0.1 and 1.0µM) and significantly lower than that in explants treated with 200μM nicotine alone (0.1 and 1.0µM; **H**). The 0.01µM MLA treatment 2 hr. prior to nicotine did not produce significant changes compared to nicotine alone (**H**). Pair-wise comparisons with Dunnett’s test (**C, E**) or Benjamini-Hochberg linear step-up procedure (**D, H**). Data represent mean±SEM.

The basal forebrain explants were cultured in Matrigel for 48 hr. in the presence of 2, 20, or 200 µM nicotine. Explants grown in culture media without nicotine served as the control. The media and nicotine were replaced approximately every 12 hrs. The nicotine concentrations used here activate nicotinic acetylcholine receptors (nAChR) in vitro ^64–66^. After 48 hrs., the explants were fixed with paraformaldehyde and migration of cells away from the explant margins was analyzed using phase contrast microscopy.

The vast majority (98%) of migrating cells were found within 720µm from the explant edge in all the experimental groups. Therefore, to minimize potential variability due to inclusion of the sparsely populated distal locations, we included in the analysis cells located within 720µm of the explant edge.

There was a significant effect of nicotine on the density of migrating neurons (*F*_(3,33)_ = 6.46, *p* < 0.01). Multiple pair-wise comparisons between each of the three nicotine doses and control explants showed a significant reduction in the number of migrating neurons for the 20µM (*t*_(33)_ = −2.95 *p* <0.05; Fig.3C) and 200µM (*t*_(33)_ = −3.12, *p* < 0.05; Fig. 3B; 3C) but not 2µM dose (*t*_(33)_ = 0.07, *p* > 0.05; Fig. 3C).

To seek direct evidence of nicotine’s effects on migratory rate, we analyzed the spatial distribution of migrating cells relative to the explant edge. For each explant, cell density was calculated at successive 80µm intervals from the explant edge up to a distance of 720µm and expressed at each interval as a percentage of the total number of cells. Higher percentage of cells nearer the explant edge and fewer cells reaching the distal intervals would serve as a proxy measure for reduced migratory rate.

There was significant interaction between the effects of nicotine and migratory distance (*F*_(24,377)_ = 6.53, *p* < 0.0001). Pair-wise comparisons revealed a significant increase in the percentage of cells located closest to the explant edge (i.e., within the initial 80µm) in all three nicotine doses compared to control [2µM > 0µM; (*t*_(377)_ = −5.70, *p* < 0.05); 20µM > 0µM; (*t*_(377)_ = −8.13, *p* < 0.05); 200µM > 0µM; (*t*_(377)_ = −10.47, *p* < 0.05; Fig. 3D)]. Thus, the migratory distance data offer compelling indirect evidence for reduced rate of migration following exposure to nicotine.

### The Effect of Nicotine on GABA neuron Migration is Mediated by the α7 nAChR

Nicotine’s biological effects are mediated by nAChRs, which are ligand-gated ion channels composed of various α and β subunits. The subunit combinations confer distinct pharmacological and physiological properties. In the developing brain, the α7 homomeric receptor is one of the most prevalent subtypes and is implicated in regulating key neurodevelopmental processes, including neuronal migration ^51,67,68^. Specifically, activation of the α7 nAChR is linked to cytoskeletal remodeling affecting neuronal migration and axonal pathfinding ^69–71^. To evaluate whether nicotine’s effects on GABA neuron migration are mediated via α7 nAChR activation, we examined receptor-specific mechanisms.

We cultured the E15 GE explants collected from exposure-naïve embryos in the presence of the selective α7 nAChR agonist, PNU282987 (0.1, 1, 10µM) for 48 hrs. The media containing the agonist was refreshed approximately every 12 hrs. We found a significant effect of PNU282987 on the density of cells migrating from the explant (*F*_(3,24)_ = 5.44, *p* < 0.05). Multiple pair-wise comparisons among the different PNU282987 doses and control showed that migration was reduced significantly following treatment with 0.1µM PNU282987 (*t*_(24)_ = −3.15, *p* < 0.05; Fig. 3B; 3E). There was no significant difference between 1µM and control (*t*_(24)_ = 0.37, *p* > 0.05; Fig. 3E) or 10 µM PNU282987 and control (*t*_(24)_ = 0.56, *p* > 0.05; Fig. 3E).

When we analyzed the spatial distribution (as percentage of the total number of cells) of migrating cells relative to the explant edge, there was no significant effect of PNU282987 (*F*_(3, 237)_ = 0.10, *p* > 0.05; Fig. 3F). The interaction between PNU282987 and intervals was also not significant (*F*_(24, 237)_ = 1.04, *p* > 0.05).

Nicotine and the α7 nAChR agonist PNU282987 significantly reduced the density of migrating cells, suggesting that nicotine’s effects may be mediated via the α7 nAChR. To determine if nicotine’s effects were mediated via the α7 nAChR, we conducted experiments using a combination of nicotine and the selective α7 nAChR antagonist methyllycaconitine (MLA). E15 GE explants collected from exposure-naïve embryos were cultured with MLA (0.01, 0.1, or 1 µM) for 2 hr. prior to the addition of 200 µM nicotine, the highest concentration that significantly reduced cell migration (Fig. 3C).

To rule out constitutive activity of α7 nAChRs, we assessed if treating GE explants with MLA in the absence of nicotine influenced GABA neuron migration. While constitutive activity (receptor activity in the absence of an agonist) of α7 nAChRs has been observed in certain experimental systems and immune cells ^72–74^, it has not been reported in embryonic GABA neurons *in vivo* or *in vitro*. We compared the density of GABA neurons treated with MLA only (0.01, 0.1, 1μM) to the control (plain media). MLA alone did not alter the density of migrating GABA neurons (*F*_(32)_ = 0.98, *p* > 0.05) supporting the absence of constitutive α7 nAChR activity.

Next, we compared the density of migrating GABA neurons in explants that were treated with MLA (0.01, 0.1, 1μM) for 2 hrs. prior to the addition of nicotine (200 µM). Explants treated with nicotine (200µM) alone or plain media (control) from previous experiments (data shown in Fig. 3C) were re-incorporated into the study design to facilitate comparisons. The effect of treatment was significant (*F*_(4,25)_=4.32, *p* < 0.05). Pair-wise comparisons showed that the density of migrating GABA neurons in explants treated with each of the 3 concentrations of MLA prior to nicotine was significantly higher than that in explants treated with nicotine alone (MLA + 200μM nicotine>200μM nicotine: 0.01μM, *t*_(25)_ = 2.57, *p* < 0.05; 0.1μM, *t*_(25)_ = 2.24, *p* < 0.05; 1μM, *t*_(25)_ = 3.66, *p* < 0.01; Fig. 3B; 3G). Compared to control explants (plain medium), GABA neuron density was not significantly different when the explants were pretreated with MLA prior to nicotine, at any concentration of MLA (control vs MLA + 200μM nicotine: 0.01μM, *t*_(25)_ = −0.26, *p* > 0.05; 0.1μM, *t*_(25)_ = 0.04, *p* > 0.05; 1μM, *t*_(191)_ = −1.25, *p* > 0.05; Fig 3G). These data show that blocking the α7 nAChR with MLA prevented the nicotine-induced reduction in the density of GABA neurons migrating from the explant suggesting a role for the α7 nAChR in mediating nicotine’s effects.

When we analyzed the percentage of migrating cells located at each of the successive 80 µm intervals from the explant edge up to a total of 720 µm, there was a significant interaction between the different treatments and distance of migratory distance (*F*_(32, 304)_ = 6.34, *p* < 0.0001). Pair-wise comparisons demonstrated consistent with the data in Fig. 3C, that nicotine produced a significant increase in the percentage of cells located within the initial 80 µm of the explant edge compared to the control (Fig. 3H). Pretreatment with MLA prior to 200 µM nicotine produced significant increases in this measurement compared to the control (MLA > control, 0.01µM, t_(304)_ = −7.16, *p* < 0.0001; 0.1µM t_(304)_ = −7.72, *p* < 0.0001, 1.0µM t_(304)_ = −6.14, *p* < 0.0001; Fig. 3H) and significant reductions compared to treatment with nicotine alone (MLA < nicotine, 0.1µM t_(304)_ = −3.49, *p* < 0.001, 1.0µM t_(304)_ = −4.54, *p* < 0.0001; Fig. 3H). Thus, although, pre-treatment with MLA blocked the effects of nicotine on neuronal migration when the density of migrating GABA neurons across the full 720 µm field was analyzed (Fig. 3G), nicotine’s effects were not blocked by MLA when migrating neuron density was analyzed at 80 µm intervals. Further research is needed to establish whether the differences between the two types of analyses are due to methodological factors alone or have biological relevance as well.

## Discussion

Our *in vivo* and *in vitro* findings collectively indicate that e-cigarette aerosol exposure is associated with significant reduction in the migration of GABA neurons from the GE to the developing cerebral cortex in the embryonic mouse brain. Furthermore, we identified nicotine—rather than the propylene glycol and vegetable glycerin mixture in the aerosol—as the active component of e-cigarette aerosol that was responsible for the effect. Moreover, our in vitro experiments revealed that the effects of nicotine are mediated via the α7 nAChR.

Our study is the first to establish a receptor-mediated mechanism by which nicotine, delivered via e-cigarette aerosol, disrupts a key developmental process in the embryonic brain, namely GABA neuron migration. Our results offer a mechanistic foundation for our earlier findings of reduced GABA neuron-to-projection neuron ratios in the adult frontal cortex following early-life nicotine exposure martin. In addition, our findings have implications for understanding the developmental origins of neuropsychiatric conditions such as ADHD, autism and schizophrenia, which have been associated with compromised GABA function and prenatal nicotine exposure ^18-20,40-43^.

Our aerosol exposure paradigm has high translational relevance. The timing of the exposure was designed to mimic nicotine use during the first trimester of human pregnancy ^75^, including the preconception period. Many women who use nicotine products may do so unknowingly during early gestation, as pregnancy often remains unrecognized until several weeks post-conception ^6,23^. Importantly, prior reports indicate that nicotine exposure during early pregnancy exerts more pronounced effects on fetal brain development than exposures occurring later in gestation smith.

Consistent with some prior studies, we did not observe adverse effects of e-cigarette or e-liquid aerosol exposure on maternal body weight gain during pregnancy, pregnancy outcomes, or gross features of embryonic development. However, published findings on these outcomes are highly variable across both human ^28,29,76^ and animal studies models ^32–34,36,37,39,77–81^. In rodents, differences in exposure timing, aerosol composition, strain background, and experimental design may contribute to inconsistent outcomes. Careful standardization and further investigation are required to clarify these discrepancies and improve the translatability of animal models.

Although we did not observe significant effects of e-liquid aerosol without nicotine on GABA neuron migration, we acknowledge that other studies have reported transcriptomic and inflammatory changes in the brain following perinatal exposure to e-liquid aerosols during brain development ^32,33^. Effects on non-neural organs such as the lungs and brain vasculature have also been described ^34,82^. These reports underscore the need for further investigation before considering nicotine-free e-cigarette aerosols safe during pregnancy.

The reduction in GABA neuron migration following prenatal e-cigarette aerosol exposure was evident at the rostral but not caudal level of the forebrain. Regional heterogeneities in transcription factor profiles of GABA neurons and their precursors may modify response to e-cigarette aerosols ^60,83,84^. The effect of e-cigarette aerosol exposure was selective to GABA neurons in the MZ of the developing dorsal forebrain. Segregations of GABA neurons into the different laminae (MZ, CP, IZ) reflects differences in their basal forebrain of origins as well as differences in cell extrinsic signals in the different laminae of the dorsal forebrain ^85^. Therefore, it is possible that e-cigarette aerosol exerts different effects on different subpopulations of GABA neurons based on cell-intrinsic and -extrinsic characteristics. Further research is warranted to explore this cell-type and region-specific vulnerability.

Endothelial cells of the embryonic forebrain emerge as likely cell-extrinsic signals that mediate region-specific effects of e-cigarette aerosol on GABA neurons. Endothelial cells express nAChR receptors ^86^ and MZ endothelial cells are transcriptionally different from IZ endothelial cells ^87^. Moreover, neurovascular effects of nicotine, combustible cigarettes and e-cigarette aerosols are well documented ^88–90^. Thus, although our prior work has demonstrated that endothelial cells can guide GABA neuron migration in the embryonic forebrain ^61,91^, whether e-cigarette aerosol exposure disrupts this guidance through direct endothelial interactions or secondary signaling pathways remains to be determined.

Our *in vitro* explant culture studies demonstrated that exposure of the GE explants to nicotine reduced the migration of GABA neurons. Furthermore, the effects of nicotine were mediated via the α7 nAChR. This was demonstrated first in explants exposed to the selective α7 nAChR agonist, PNU282987 and in explants exposed to the selective α7 nAChR antagonist MLA prior to exposure to nicotine.

We did not perform real-time live-cell imaging of neuronal migration in our explant cultures. Therefore, our findings are not derived from a direct measurement of neuronal migration in real time. However, the reduction in GABA neuron density in the dorsal forebrain in the in vivo studies, reduction in the density of cells exiting the explant following exposure to nicotine and increased accumulation of neurons near the explant periphery (~80 µm from the edge) in the nicotine-exposed explants collectively indicate reduced GABA neuron migration. Our data cannot rule out neurogenesis ^92^ or cell death ^93^ as contributing factors.

Our in vitro data suggest that the effects of nicotine and the α7 nAChR agonist are dose-dependent and are unlikely to be toxic effects. In addition, the accumulation of migrating neurons near the explant margins shows effects on migration independent of potential effects on cell death. However, direct evaluation of neurogenesis and cell death would be needed to firmly rule out the potential contribution of these processes to our findings.

The effects of the α7 nAChR agonist PNU282987 on neuronal migration *in vitro* were observed at a concentration of 0.1 µM but not at higher concentrations (1 or 10 µM). The lack of effect at higher concentrations is consistent with known properties of the α7 nAChR, which rapidly desensitizes and inactivates upon sustained or repeated stimulation, which can occur at elevated ligand concentrations ^72,94^. Our explant experiments using the selective α7 nAChR antagonist MLA further suggest that additional nAChR subtypes may contribute to nicotine’s effects on GABAergic neuron migration. Specifically, while MLA pretreatment blocked nicotine’s effects when migration was quantified by the density of neurons migrating from the explant (Fig. 3G), it did not fully block nicotine’s effects when migration was assessed at 80 µm intervals (Fig. 3H) pointing to the likely involvement of other nAChR subtypes, other than α7 nAChR. Further research is warranted to elucidate the specific receptor subtypes and signaling pathways involved. Our findings pave the way for future research on this topic.

In summary, our study provides novel evidence that prenatal e-cigarette aerosol exposure reduces GABA neuron migration in the developing forebrain. These effects are attributable, at least in part, to nicotine acting via α7 nAChRs. Our findings uncover a previously unrecognized vulnerability of GABA neurons of cerebral cortical circuits to nicotine and highlight a potential mechanistic link between prenatal nicotine exposure and neurodevelopmental disorders. Furthermore, our findings underscore the urgent need for updated educational campaigns, clinical guidelines, and policy interventions that explicitly address the neurodevelopmental risks of e-cigarettes. Without such measures, the use of e-cigarettes during pregnancy may contribute to a silent burden of neurodevelopmental disease risk for the next generation.

## Methods

### Animals

Swiss Webster (SW) mice (Charles River Laboratories, Wilmington, MA) and GAD67-GFP^+/−^ knock-in mice ^54^ were housed at the institutional laboratory animal housing facility in individually ventilated, temperature-, and humidity-controlled cages on a 12/12hr light/dark cycle. Water and food were provided *ad libitum*. This study protocol was reviewed and approved by the Florida State University Animal Care and Use Committee. All experiments complied with NIH guidelines for the care and use of laboratory animals.

### Aerosol Exposure

The exposures were conducted using the Buxco E-cigarette/Vapor/Tobacco Smoke Generator system (Data Sciences International, St Paul, MN) equipped with three 9L exposure chambers for whole-body aerosol exposure. Each chamber held 8 mice, each in separate “cubicles”. A separate chamber was used for each exposure to avoid cross-contamination of the equipment by the aerosols. Female mice were randomly assigned to one of three exposure groups: e-cigarette aerosol, e-liquid aerosol, and room air. Mice in all three exposure groups were habituated to ambient room air (flow rate: 4L/min) in the exposure system for 48 or 96 min per day for 5 days. Following habituation, each group was acclimated to their respective exposure conditions for 48 or 96 min/day, 5 days/week for 2 weeks (Fig. 1).

The e-cigarette devices (Smok X-Priv kits, Vapor Authority, San Diego, CA) were used at 100W with a 0.12Ω coil (Vapor Authority, San Diego, CA). E-liquid (PG:VG = 50:50) with 2.4% nicotine and without nicotine. (Vapor Vapes, Sand City, CA). The e-cigarette and e-liquid aerosol were delivered in 55mL puffs created with a 2-second inhale and 4-second exhale and delivered at a rate of 1 puff/min ^95^.

After acclimation, the female mice were paired with unexposed GAD67-GFP^+/−^ sires during the night (dark cycle) and checked for the presence of a vaginal plug in the morning. The pairing occurred every night until a plug was discovered or for a maximum of 8 nights (to include two estrous cycles). The presence of a vaginal plug was recorded as gestational day zero (GD0). The exposure continued every day up to and including GD14 (Fig. 1)

### Cotinine Analyses

Dams exposed to e-cigarette aerosol for 48 or 96 minutes (n=3-4) were anesthetized with isoflurane and decapitated. Trunk blood samples were collected and following 15 min at room temperature, the samples were centrifuged for 10 minutes at 3000 rpm. Serum was collected as the supernatant, frozen in liquid nitrogen and stored at −80°C until further analysis.

Embryos (E14) were removed from anesthetized dams (ketamine 90mg/kg and xylazine 10mg/kg; i.p.) via hysterectomy and the dams were euthanized by exsanguination. Forebrains were removed, lysed in RIPA buffer (50 mM Tris-Cl pH 8.0, 150 mM NaCl, 2 mM EDTA, 1% NP-40, 0.1% SDS, 10 mM NaF) with 1% aprotinin (Affymetrix #101172-198), leupeptin (Alfa Aesar #AAJ61188-MC) and 1% AESBF (Alfa Aesar #AAH26473-MD) protease inhibitors. Protein in the lysates was quantified using the Qubit Protein Assay (ThermoFisher #Q33211) and Qubit 2.0 Fluorometer. Lysates were frozen in liquid nitrogen and stored at −80°C until further analysis.

Serum and brain cotinine content were analyzed with ELISA (Immunoalysis, #217-0096). Approximately 10 µl of serum and 330 µg of protein from E14 brain samples were loaded in duplicate. The serum cotinine concentration was averaged from 3-4 dams. Brain cotinine concentration from 2-3 embryos per litter was averaged and litter averages (3-4 litters) used to estimate E14 brain cotinine concentration.

### Collection of Embryonic Brains for Histological Analyses

On GD15 the dams were anesthetized [ketamine (90mg/kg) and xylazine (10mg/kg); i.p.] and the embryos were removed by hysterectomy. GAD67-GFP^+/−^ embryos were identified using fluorescence goggles (Biological Laboratory Equipment, Budapest, HU). GFP+ embryonic brains were dissected, and immersion fixed with 4% paraformaldehyde at 4°C overnight. Brains were subsequently cryoprotected with 15% sucrose for 24 hrs. followed by 30% sucrose for 24 hrs. at 4°C. The cryoprotected brains were flash frozen on dry ice and stored at −80°C. Deeply anesthetized dams were euthanized by exsanguination.

### Analysis of GABA Neuron Density in the E15 Dorsal Forebrain

The brains were cut into 20 µm-thick sections in the coronal plane using a cryostat and mounted on SuperFrost Plus slides (Fischer Scientific, #12-550-15; 2 embryos from each of 4 litters). Two rostral sections and two caudal sections per brain were chosen based on anatomical landmarks and imaged with the Keyence BZ-X700 microscope. The two sections from each rostro-caudal level were approximately 340 μm apart. Rostral sections were at the coronal plane where the lateral and medial ganglionic eminences separated by the sulcus were visible (Fig. 2A). Caudal sections were taken at the coronal plane where only the caudal ganglionic eminence was visible (Fig. 2E). The caudal sections were approximately 360 µm caudal to the rostral sections. Average values of the two rostral and two caudal sections were calculated for each embryo.

The number of GAD67-GFP+ cells were counted within a grid superimposed digitally on the image of the dorsal cerebral wall. The grid was 100 µm wide and its height encompassed the full radial thickness of the dorsal cerebral wall from the ventricular to the pial surface (Fig. 2A, E). The grid was divided into 10 equal-sized “bins” along the radial axis extending from the ventricular to pial surface. GFP+ cells were counted within each “bin”. This method allowed us to count the cells and record their position along the radial axis of the dorsal cerebral wall so that each cell could be assigned its position within the MZ, CP, IZ and SVZ/VZ. We have used this counting method previously^53^.

The grids were superimposed at each of 3 locations along the dorsal cerebral wall namely, lateral, middle, and medial anatomical locations. The lateral location was approximately 100 µm dorsal to the caudate-pallial angle (Fig. 2A, E). The middle location was 100 µm lateral to the interhemispheric sulcus and the medial location was in the dorsal forebrain that is adjacent to the interhemispheric sulcus. The locations were chosen using the same criteria at the caudal level as well. The three locations permitted analysis of the data at multiple positions along the lateral to medial maturational gradient.

### Basal Forebrain Explant Cultures

Exposure naïve SW females and males were bred to produce timed pregnancies. Pregnant dams were anesthetized [ketamine (90mg/kg) and xylazine (10mg/kg); i.p.] on GD15 and the embryos were removed by hysterectomy. The brains were dissected, and the forebrain was embedded in low melting point agarose (4%; Type VII-A; Sigma Aldrich, #A0701). The forebrains were cut in the coronal plane using a Vibratome into 250 µm-thick sections. The sections were collected and stored in 1X Krebs buffer (NaCl 126mM, KCl 2.5mM, NaH_2_PO_4_ 1.2mM, MgCl_2_ 1.2mM, CaCl_2_ 2.5mM) at 4°C for up to 2 hours. The sections were incubated in fetal calf serum (ThermoFisher, #10082147) and MEM Eagle’s medium (Sigma Aldrich M-4780) supplemented with 45% D-(+)-Glucose (Sigma Aldrich, G-8769), 100X pen/strep (Sigma Aldrich, #P0781) with glutamine (Sigma Aldrich, #G7513) for 30 minutes at 37°C and 5% CO_2_.

Coronal sections representing the rostral forebrain were identified using the same landmarks that were used for the *in vivo* studies. From each rostral section, bilateral basal explants were collected using a 1mm biopsy punch (Fisher Scientific, #NC9226138). The 1mm explant included the MGE and the developing striatum (Fig. 3A). The use of 250 µm thick sections and the biopsy punch ensured uniformity of size (diameter and thickness) of the explants across embryos. Each explant was plated in 50µL of a 50:50 mixture of Matrigel matrix (Fisher Scientific, #p-90960) and Eagles MEM medium in a clear bottom, 96-well plate. The Matrigel was allowed to set for 30-minute at 37°C and 5% CO_2_. Serum-free Neurobasal Medium (Gibco, #21103-049) supplemented with B27 (Gibco, #17504-044), 45% D-(+)-Glucose, 100X pen/strep/glutamine was added to the wells and the explants were incubated at 37°C and 5% CO_2_ for 48 hrs.

Following 48 hr. in culture, the medium was replaced with 4% PFA for 10 min to fix the explants still encased in the Matrigel matrix, and washed 3 times, 5 minutes each with PBS. Phase contrast images of cell migration were acquired using the Keyence BZ-X800 microscope.

### Drugs

Nicotine (Sigma Aldrich #N3876), the selective α7 nAChR agonist PNU282987 hydrate (Sigma Aldrich #P6499) and the selective α7 nAChR antagonist methyllycaconitine (MLA; MedChemExpress LLC #502261192) were used. PNU282976 was dissolved in 0.1% DMSO (Sigma Aldrich #D8418) and MLA solid was dissolved in dH_2_0. The timing of drug application and replenishment are described in the Results section.

### Analysis of neuronal migration

A sampling method modified from our previous study was used to achieve unbiased sampling of cell migration^56^. Only those explants that exhibited cell migration from at least 50% of the periphery were chosen for analysis. Images of the explant and the migration fields were collected, and a rectangular grid (200 µm wide and 720 μm in length) was digitally superimposed on the image. The grid was positioned such that the base of the grid was parallel to the edge of the explant (Fig. 3B). Two grids were superimposed on images of each explant. The position of the first grid was selected by the experimenter so that it overlaid robust migration, and the second grid was positioned to be 90° clockwise from the first regardless of the quality of the migration field, to achieve unbiased sampling. The long axis of the grid was divided into 80 µm “bins”. GABA neuron density was calculated by dividing the total number of migrating cells by the area of the sampling region. For each explant, GABA neuron density was calculated as the average of data from two grids. Two to 3 explants were used per embryo. One to 3 embryos were used per litter. For each experimental group 3-4 litters were used.

### Statistical Analyses

Maternal bodyweight, litter size, crown-rump length, GABA neuron density, and migratory distance were analyzed using general mixed model framework (random effects ANOVA analyses with litter as a random effect, and mixed model ANOVAs; with between and within subject factors. Specifically, in vivo and in vitro GABA neuron density and radial thickness of the dorsal cerebral wall were analyzed using a mixed-model ANOVA to evaluate between-subject and within-subject factors. Between-subject factors were treatment (*in vivo*: Room air, E-liquid, E-cigarette; *in vitro*: nicotine, PNU282987, MLA). Anatomical location (lateral, middle, and medial) and migratory zone (MZ, IZ, CP and SVZ/VZ) were treated as fixed factors. Migratory distance (80 µm intervals) was a within-subject factor. Litter was treated as a random factor. Embryo was an additional repeated measure within-subjects factor for the in vivo analyses of GABA neuron density. Statistically significant (*p*⍰< ⍰0.05) findings were followed up by simple contrasts tests to evaluate any significant between-subject and within-subject effects and their interactions (i.e., interactions among treatment, anatomical location, and migratory zone). Pair-wise comparisons were used to directly compare statistically significant main effects to controls, applying Dunnett’s test for multiple comparisons. Pair-wise comparisons using Bonferroni’s tests were conducted in instances where a control group is not defined such as in the analyses of cell density and radial height between anatomical locations. We controlled for type I error in the post hoc analysis of those significant interactions using the linear step-up procedure. The analyses were performed using SAS/STAT 9.4 (SAS, Cary, NC). Graphs were prepared using Prism 10.1.2 (GraphPad Prism, San Diego, CA).

The study was designed, performed and is reported in accordance with ARRIVE guidelines (https://arriveguidelines.org).

## Data Availability Statement

All data generated or analyzed during this study are included within the article.

